# Contrastive Representation Learning for Single Cell Phenotyping in Whole Slide Imaging of Enrichment-free Liquid Biopsy

**DOI:** 10.1101/2025.05.21.655334

**Authors:** Amin Naghdloo, Dean Tessone, Rajiv M. Nagaraju, Brian Zhang, Jeffrey Kang, Shouyi Li, Assad Oberai, James B. Hicks, Peter Kuhn

## Abstract

Tumor-associated cells derived from a liquid biopsy are promising biomarkers for cancer detection, diagnosis, prognosis, and monitoring. However, their rarity, heterogeneity and plasticity make precise identification and biological characterization challenging for clinical utility. Enrichment-free approaches using whole slide imaging of all circulating cells offer a comprehensive and unbiased strategy for capturing the full spectrum of tumor-associated cell phenotypes. However, current analysis methods often depend on engineered features and manual expert review, making them sensitive to technical variations and subjective biases. These limitations highlight the need for a better feature representation to improve performance and reproducibility of applications in large-scale patient cohort analyses. In this study, we present a deep contrastive learning framework for learning features of all circulating cells, enabling robust identification and stratification of single cells in whole slide immunofluorescence microscopy images. We demonstrate performance of learned features in classification of diverse cell phenotypes in the liquid biopsy, achieving an accuracy of 92.64%. We further demonstrate that learned features improve performance in downstream applications such as outlier detection and clustering. Lastly, our feature representation enables automated identification and enumeration of distinct rare cell phenotypes, achieving average F1-score of 0.93 across cell lines mimicking circulating tumor cells and endothelial cells in contrived samples and average F1-score of 0.858 across CTC phenotypes in clinical samples. This workflow has significant implications for scalable analysis of tumor-associated cellular biomarkers in clinical prognosis and personalized treatment strategies.

## 1 Introduction

Liquid Biopsy (LBx) offers a non-invasive method for cancer detection, progression monitoring, and assessment of minimal residual disease by sampling biomarkers from bodily fluids using techniques such as flow cytometry and whole slide imaging (WSI) [1–5]. LBx has advantages compared to traditional tissue biopsy by enabling non-invasive repeated sampling to obtain longitudinal molecular information from tumors [4]. Circulating tumor cells (CTCs), malignant cells disseminated from the tumor into the peripheral bloodstream capable of seeding distant metastatic lesions, are the primary cells studied in LBx [6, 7]. Elevated CTC counts in patients are associated with poorer clinical outcomes, including reduced progression-free survival and overall survival, across multiple cancer types [7–11]. CTCs exhibit significant plasticity and heterogeneity and have been categorized into several phenotypes associated with poor patient outcomes, including epithelial-to-mesenchymal transition (EMT) [12, 13], homogeneous and heterogeneous cell clusters [14], platelet-coated CTCs [15], and immune-like (CD45+) CTCs [16]. Additionally, previous studies have identified and investigated the utility of other tumor-associated cell populations in peripheral blood [8], such as circulating endothelial cells [17], cancer-associated fibroblasts [18], and macrophages [19]. Recent studies have also demonstrated the prognostic value of immune cell subpopulations in patients with solid tumors [20]. It was shown that leukocytes within heterogeneous CTC clusters may inform tumor–immune interactions and patient stratification [21], underscoring the value of profiling immune phenotypes in liquid biopsy as well. The study of tumor-associated cell types and phenotypes can provide a more comprehensive view of tumor heterogeneity and evolution in systemic circulation, underscoring the need for approaches that describe their full diversity.

One of the main challenges in cell-based LBx technologies is the extreme rarity of circulating tumor-associated cells, often occurring at rates below one per million leukocytes [22]. These cells are typically studied using biophysical enrichment platforms [23], which enhance sensitivity by targeting predefined phenotypic traits such as cell size or surface marker expression to isolate specific cell populations. Recent developments in deep learning methods have substantially improved the detection of CTCs, facilitating more accurate and rapid identification [24]. While effective for detecting canonical CTCs, this targeted approach reduces the ability to capture the full heterogeneity of the circulating cell population. Alternatively to biophysical enrichment, enrichment-free platforms seek to profile all cells by plating them on slides and using immunofluorescent (IF) markers to differentiate cell phenotypes, generating WSI data that enable comprehensive profiling of the heterogeneous population of tumor-associated cells among all cells [25, 26].

Current analytical methods for LBx WSI data largely rely on engineered features [15, 27–29], yet selecting these features demands substantial apriori knowledge of and depends on the cell phenotypes under study. After feature selection, these features are used to detect candidate rare cells, which are then manually reviewed, curated and phenotypically classified by human experts. However, engineered features are sensitive to technical variations common in whole slide imaging, such as blurry regions, which can introduce undesirable variance to downstream analyses [30–32]. Moreover, expert-driven annotation and classification are susceptible to subjective bias and lack scalability, limiting throughput and reproducibility of large scale analyses across patient cohorts. These limitations highlight the necessity of data-driven workflows to extract robust features from cell images in order to enhance algorithmic methods for identification and phenotypic characterization of tumor-associated rare cells, thereby minimizing human involvement in the process and improving scalability in large scale studies.

Deep representation learning has emerged as an effective approach for learning robust features from image data [33–35]. These models are often trained in a self-supervised manner, allowing them to learn unbiased representations of cell images without the need for extensive labels generated by humans [36, 37]. Among representation learning techniques, contrastive learning has been shown to learn powerful discriminative representations across diverse natural images [38]. Several studies in drug discovery have applied contrastive learning to single cell IF microscopy images, where it enables mechanism-of-action classification through single cell representations [39–41]. While these studies demonstrate the utility of self-supervised learning methods for prevalent cell phenotypes, their potential for representing the rare and heterogeneous cell phenotypes present in LBx remains largely underexplored.

In this study, we describe a deep learning framework for extracting robust features from single cell images in LBx that enables resolving the entire phenotypic heterogeneity of the rare tumor-associated cells in WSI data. The approach comprises two main modules: a segmentation model and a feature extraction model. We trained and validated both components using curated well-balanced datasets from patient samples to better represent the rare cell phenotypes. We further demonstrate the utility of the learned features in a series of downstream tasks, including classification of diverse cell phenotypes, outlier detection for identifying rare cell phenotypes, clustering distinct phenotypes, and finally the application of identification and enumeration of rare cell phenotypes in WSI data with severe imbalance using both cell line spiked samples and patient samples. Together, this work provides a framework to learn a robust feature space enabling a highly scalable clinical tool for accurate identification and enumeration of cell biomarkers in LBx and underpinning the unsupervised identification of novel cell phenotypes in WSI data.

## 2 Results

### 2.1 Overview of the single cell phenotyping framework

Here we establish a deep learning framework designed for robust feature extraction from single cells in WSI data to enhance the performance of common downstream tasks in LBx data analysis (Figure 1a). Our framework leverages two primary trained modules: a segmentation model and a feature extraction model. The training data for both of these models included 406 WSI data collected from a rich dataset of 277 human-annotated patient samples from previous studies covering a heterogeneous spectrum of rare cell phenotypes identified and enumerated in patient cohort studies. The break-down and use of datasets can be found in the Methods section and Supplementary Tables 1 and 2. These WSI data were prepared using a four-color immunofluorescence assay described in detail in the Methods section. Briefly, nucleated cells were plated as a monolayer on glass slides and were stained with DAPI (for DNA), a cocktail of cytokeratins (for epithelial cells), vimentin (for mesenchymal cells), and CD45 and CD31 (for leukocytes and endothelial cells, respectively). Figure 1b represents a gallery of diverse cell phenotypes from the training datasets in this study.

**Fig. 1.**
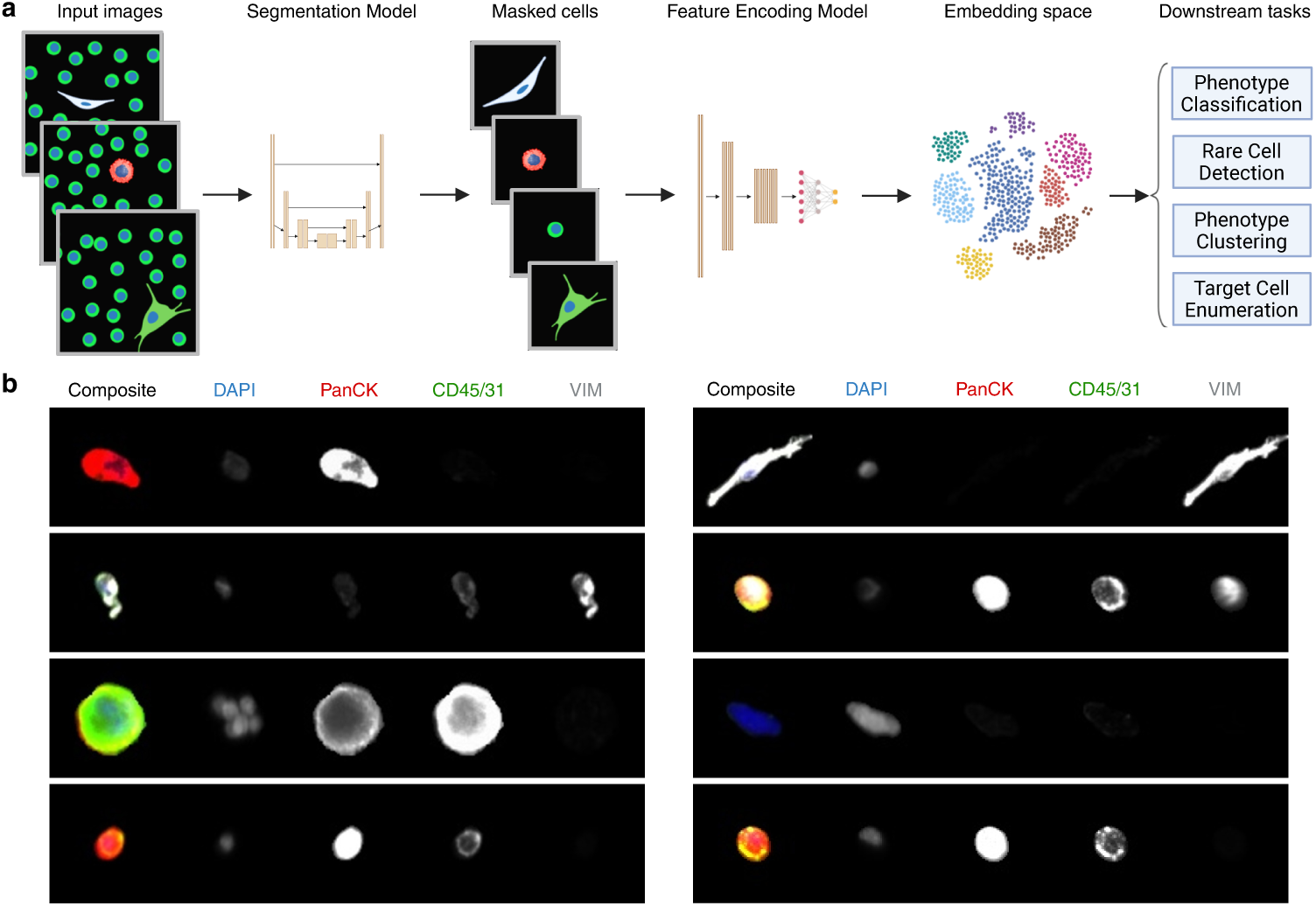
Deep phenotyping framework enables resolving heterogeneity of cell phenotypes in WSI data from LBx samples. a. Schematic overview of the deep phenotyping framework. WSI data, stored as individual input images, are first segmented to identify individual cells and their masks. Individual cell images are passed to the feature encoding module, an autoencoder trained on cell image data via contrastive learning, to generate cell image features. The encoded features can then be used for downstream tasks including classification, rare cell detection, and enumeration of target cell phenotypes for biological or clinical applications. b. Gallery of diverse cell phenotypes present among rare cells of LBx samples. A scale bar of 10 microns is shown in the bottom right.

The segmentation model, based on a U-Net architecture provided by Cellpose [42], was trained on a set of 634 large and annotated image crops, each containing at least one rare cell. Testing the model on a separate set of 158 image crops showed improved accuracy in detecting individual cells compared to the general-purpose Cellpose model. We evaluated performance using the F1-score across different thresholds of intersection-over-union (IoU), a metric that measures how well predicted cell masks overlap with ground truth masks. The trained model achieved higher F1-scores for IoU values between 0.5 and 0.8, indicating better agreement in object-level segmentation, and showed comparable performance at other IoU thresholds (Supplementary Figure 1). Further details can be found in the Methods section. The feature extraction module was then trained using 103,245 segmented and sampled rare cells and leukocytes from WSI data of 25 patients (Supplementary Figure 2). Leukocytes were controllably depleted with a pretrained binary classifier that achieved 99.37% accuracy, thereby creating a balanced dataset for training the feature encoding model. We evaluated the quality and performance of learned features in a number of downstream tasks, including classification of single cell phenotypes, rare cell detection, clustering to characterize unknown cell phenotypes, and enumeration of known tumor-associated cells in WSI data both in cell line-derived experimental samples and patient samples. The details of model architectures, their training data and implementations are described in the Methods section.

### 2.2 Linear classification confirms discriminative capacity of learned features on a broad range of cell phenotypes

Deconvolving the heterogeneous phenotypes of rare tumor-associated cells as well as immune cells is essential for scalable clinical analysis and for associating distinct cell populations with clinical outcomes, leading to biomarkers discovery in preclinical settings. In order to evaluate the performance of our learned features in distinguishing cell phenotypes, we assembled a ground truth dataset comprising a heterogeneous population of cell phenotypes, verified by a single cell targeted proteomics analysis through imaging mass cytometry (IMC) and augmented with phenotypically matched cells collected from additional samples. This dataset comprises both rare tumor-associated cells and diverse leukocyte subpopulations, annotated using additional proteomic markers and distinct morphological features.

The ground truth dataset covered seven distinct rare cell phenotypes; canonical epithelial CTCs [43], immune-like CTCs (imCTCs) [16], platelet-coated CTCs (pcCTCs) [15], circulating endothelial cells (CECs) [43], megakaryocyte-like and fibroblast-like cells, and morphologically abnormal nuclei with no other IF marker expression (Large Nuclei). The megakaryocyte-like and fibroblast-like cells are cell groups that exhibit IMC marker profiles consistent with megakaryocytes and fibroblasts, respectively. We further expanded the ground truth dataset of cell phenotypes by adding cells from three major immune cell subclasses including lymphocytes, monocytes/macrophages, and granulocytes. Table 1 represents all cell phenotypes in immunofluorescent images along with their additional proteomic markers used to describe their phenotype.

**Table 1.** Description of cell phenotypes in ground truth dataset.

| Cell phenotype | Immunofluorescent markers | Additional IMC markers |
| --- | --- | --- |
| CTC | DAPI+ CK+ | CK8/CK18+ |
| imCTC | DAPI+ CK+ CD45/31+ | CK8/CK18+ CD45+ CD31- |
| pcCTC | DAPI+ CK+ CD45/31+ | CK8/CK18+ CD45- CD31+ |
| CEC | DAPI+ VIM+ CD45/31+ | CD45- CD31+ |
| Megakaryocyte | DAPI+ CD45/31+ | CD31+ CD61+ CD68+ |
| Fibroblast | DAPI+VIM+ | CD31- CD34+ CD44+ Fibronectin+ |
| Large Nucleus | DAPI+ | N/A |
| Lymphocyte | DAPI+ CD45/31+ | CD3+ or CD20+ or CD56+ |
| Monocyte/Macrophage | DAPI+ CD45/31+ | CD3- CD20- CD56- (CD14+ or CD68+) |
| Granulocyte | DAPI+ CD45/31+ | CD3- CD20- CD56- CD14- CD68- |

We used our framework to collect learned feature representations of this dataset. A two-dimensional UMAP projection of these cell features is presented in Figure 2a. We then trained a logistic regression classifier on 80% of the dataset to predict cell phenotypes. When tested on the remaining 20% of the dataset, the model achieved an accuracy of 92.64% in classifying cell phenotypes. The micro-average area under the precision-recall curve was 0.969 and the macro-average was 0.961 (Figure 2b). We further assessed the area under the receiver-operating characteristic (ROC) curve, which achieved a micro-average of 0.996 and a macro-average of 0.994. These results are comparable to those obtained with engineered features (Supplementary Figure 3), reflecting on the high classification fidelity of learned features.

**Fig. 2.**
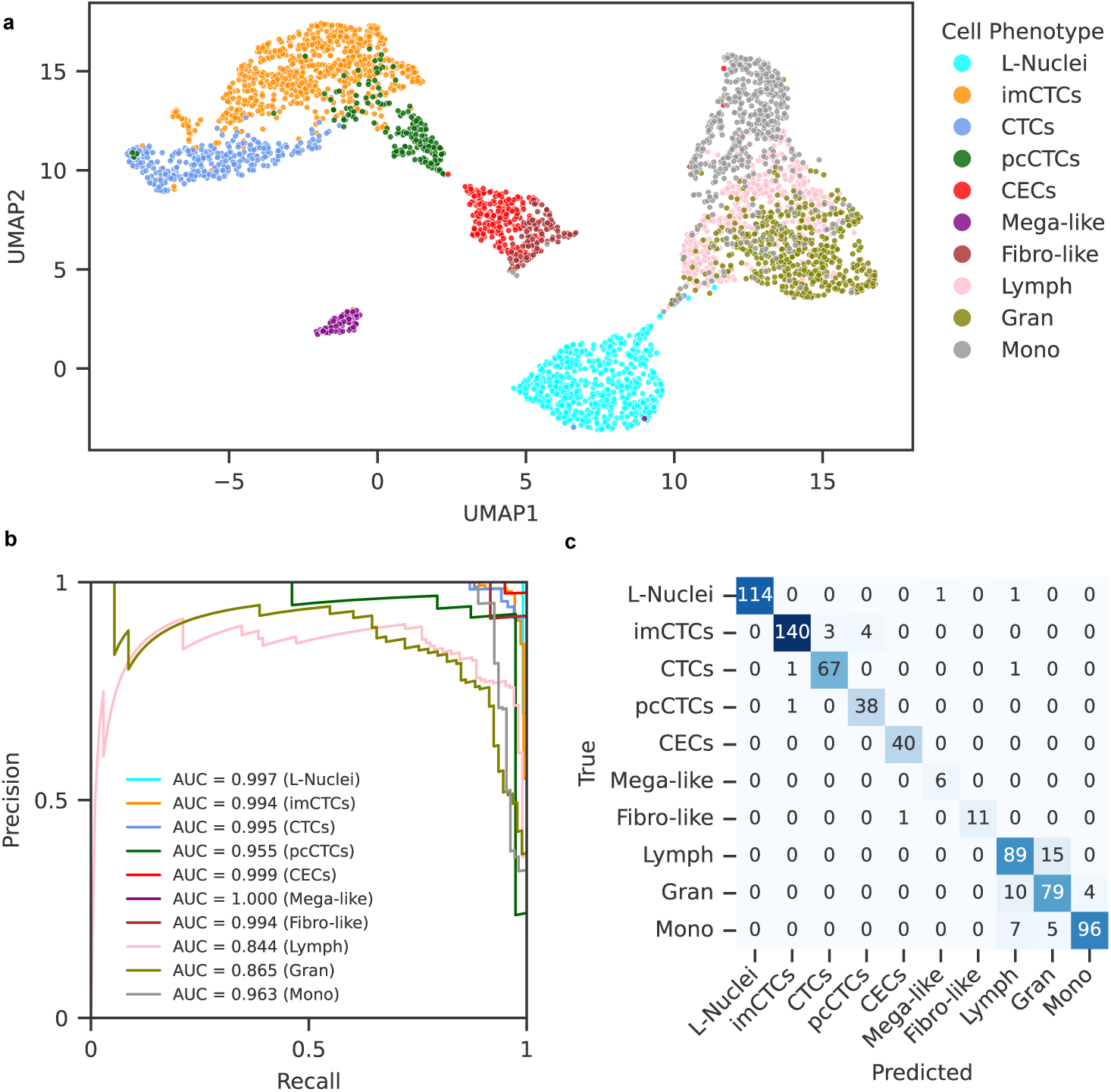
Learned feature representations stratify diverse single cell phenotypes. A ground truth dataset of diverse cell phenotypes was assembled to evaluate the performance of features provided by the deep phenotyping framework. a. UMAP projection of cells, colored by ground truth cell type annotations. b. Precision-recall curves for each cell type on the held-out test set, with areas under the curve (AUCs) indicated in the legend. c. Confusion matrix of the logistic regression multi-class classifier on the test data.

Among all rare cell phenotypes, the lowest area under the precision-recall curve is 0.955 (pcCTCs), whereas all other rare event classes have areas under the curves greater than 0.99. In addition, all classes have areas under the ROC curve of greater than 0.98. Further, we noted that the majority of misclassifications within the rare event classes naturally occur between cell phenotypes with subtle differences in their IF images (Figure 2c). Among the 11 misclassified rare events, 7 imCTCs are misclassified as either canonical epithelial CTCs (3) or pcCTCs (4). Both pcCTCs and imCTCs are epithelial cells and share identical IF identification criteria (Table 1), making them exceedingly challenging for humans to distinguish in the absence of proteomic data.

### 2.3 Contrastive learning enables inferring blur-invariant features

Technical variations in WSI, including regional blur, can compromise the accuracy of features derived from single-cell images, adversely affecting downstream phenotype identification by human analysts and, ultimately, accuracy of end results. In an attempt to mitigate these effects, we incorporated specific blur augmentations into our contrastive learning framework, resulting in a feature space robust to such technical variability, the full details of which can be found in the Methods section.

To quantify the susceptibility of single-cell features to blur, we interrogated the ground truth dataset of cell phenotypes. We applied gaussian blur with increasing standard deviation to each cell image followed by segmentation and extraction of both engineered and learned features. We then calculated the cosine distances between the original image features and their blurred counterparts. As shown in Table 2 and Supplementary Figure 4, distances in the engineered feature space are, on average, greater than those in the learned feature space, indicating a higher robustness of learned features to blurring.

**Table 2.** Cosine distance (mean ± standard deviation) between original, sharp images and blurred images for each feature space.

| Blur ($\sigma$) | Learned Features | Engineered Features |
| --- | --- | --- |
| 0.50 | $0.048 \pm 0.041$ | $0.074 \pm 0.079$ |
| 0.75 | $0.097 \pm 0.064$ | $0.195 \pm 0.128$ |
| 1.00 | $0.124 \pm 0.075$ | $0.256 \pm 0.145$ |
| 1.25 | $0.138 \pm 0.081$ | $0.296 \pm 0.152$ |
| 1.50 | $0.145 \pm 0.082$ | $0.326 \pm 0.156$ |

### 2.4 Learned features enhance outlier detection of rare tumor-associated cell phenotypes in WSI

Outlier detection is a common downstream application in enrichment-free LBx to overcome the rarity limitation of tumor-associated cells primarily in peripheral blood while preserving the cellular heterogeneity. However, the effectiveness of this approach varies across rare cell phenotypes due to differences in how these cells are represented in the feature space relative to common leukocytes. Here, we assess the performance of our learned features in detecting outliers and compare their performance to engineered features.

As a benchmark dataset to evaluate the effect of feature sets in outlier detection, we generated contrived WSI data by spiking SK-BR-3 or HPAEC or both cells into healthy blood samples at concentrations of 1:10,000 leukocytes (0.01%). SKBR3 is a human breast cancer cell line that mimics canonical epithelial CTCs, whereas, HPAECs is a human pulmonary artery endothelial cell line that mimics tumor microenvironment CECs. Slides were generated from 3 independent normal blood samples in 3 different conditions, resulting in a total of 9 slides. The three conditions were 1) normal blood spiked with SK-BR-3, 2) normal blood spiked with HPAEC, and 3) normal blood spiked with both cell lines in equal proportions (referred to as Mixed). All slides were stained and imaged with the same protocol as described in Methods, generating 9 WSI data. This dataset was processed via our trained segmentation module and manually annotated as SK-BR-3, HPAEC, or other based on marker expression. We tested both learned features extracted from our framework and engineered features to compare their effect on the performance of the outlier detection task.

We evaluated three distinct outlier detection methods on the feature sets: Copula-Based Outlier Detection (COPOD), Empirical-Cumulative-distribution-based Outlier Detection (ECOD), and isolation forest (iForest) [44]. In each test, we assessed the number of identified SK-BR-3 and HPAEC cells among the top 2,500 outliers from a total of about 2.5 million cells ( 0.1% of all cells) identified in each WSI data (Figure 3). We observed that across all three outlier detection methods and two phenotypes, the learned feature space resulted in a higher relative frequency of target cells among the identified rare events compared to engineered features. This trend was consistent across methods, indicating reproducibility of performance gains from learned features. For SK-BR-3 cells, ECOD yielded the highest area under the mean ROC curve of 0.954 utilizing the learned features, as compared to 0.517 for the engineered features. HPAEC cells were best detected utilizing COPOD with an area under the mean ROC curve of 0.938, as opposed to 0.811 for engineered features. Additionally, the detection rates across the two cell types, SK-BR-3 and HPAEC, were comparable when using the learned features. In contrast, the engineered feature space exhibited uneven detection performance, with substantially better recovery of HPAEC cells than SK-BR-3 cells.

**Fig. 3.**
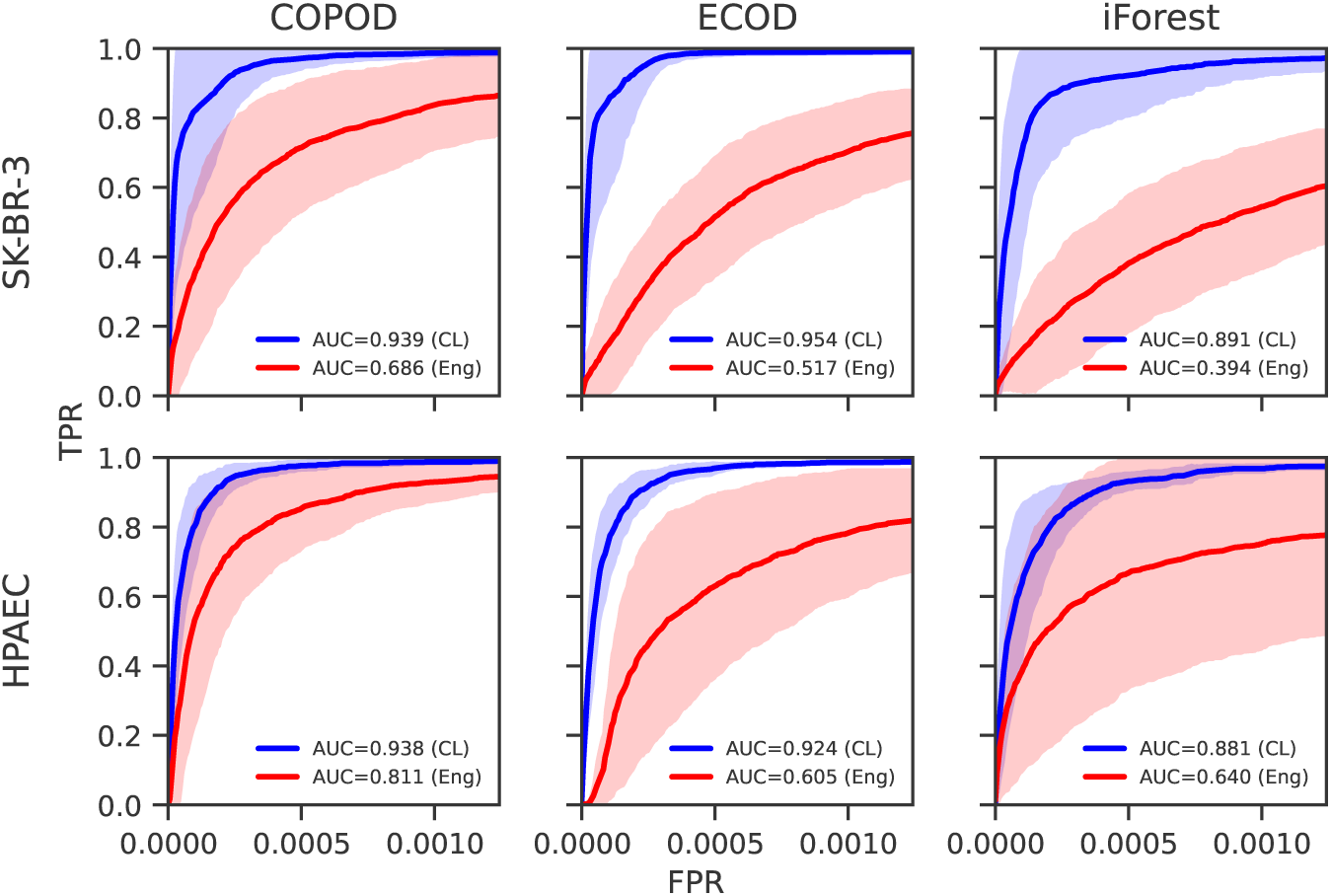
Learned feature representations enhance identification of rare cell phenotypes via outlier detection. Three outlier detection methods (COPOD, ECOD, iForest) were applied to identify rare tumor-associated cells using either learned (CL) or engineered (Eng) feature representations. The relative frequency of target cells among detected outliers (top 2500 outliers) is shown for each method and feature space.

### 2.5 Learned features improve clustering performance in presence of data imbalance

While rare cell detection methods are employed to address the challenge of extreme imbalance between immune and tumor-associated cells, the identified rare cells can still be imbalanced across phenotypes. This data imbalance can hinder downstream analyses, particularly unsupervised clustering as a common technique when investigating novel rare cell phenotypes for biomarker discovery. Clustering methods are often sensitive to data imbalance that may exist among rare cell phenotypes. While development of advanced clustering algorithms aims to address this limitation to some extent, the characteristics of the feature space on which the clustering is performed can help improve the impact of imbalance on clustering results. Here, we assess the performance of our learned features in clustering under imbalanced conditions compared to engineered features.

Leveraging the ground truth dataset of cell phenotypes, we evaluated the performance of two commonly used clustering methods, K-means and Leiden community detection, in delineating distinct cell phenotypes using learned features and engineered features. We defined the data imbalance ratio as the ratio of total immune cells to total rare cells and performed the experiment across a range of ratios from 0.5 to 10. We quantified the performance of each clustering result by calculating standard metrics of adjusted-rand index (ARI), normalized mutual information (NMI), homogeneity, and completeness (Figure 4). The description of evaluation metrics and hyperparameters of the clustering algorithms can be found in the Methods section.

**Fig. 4.**
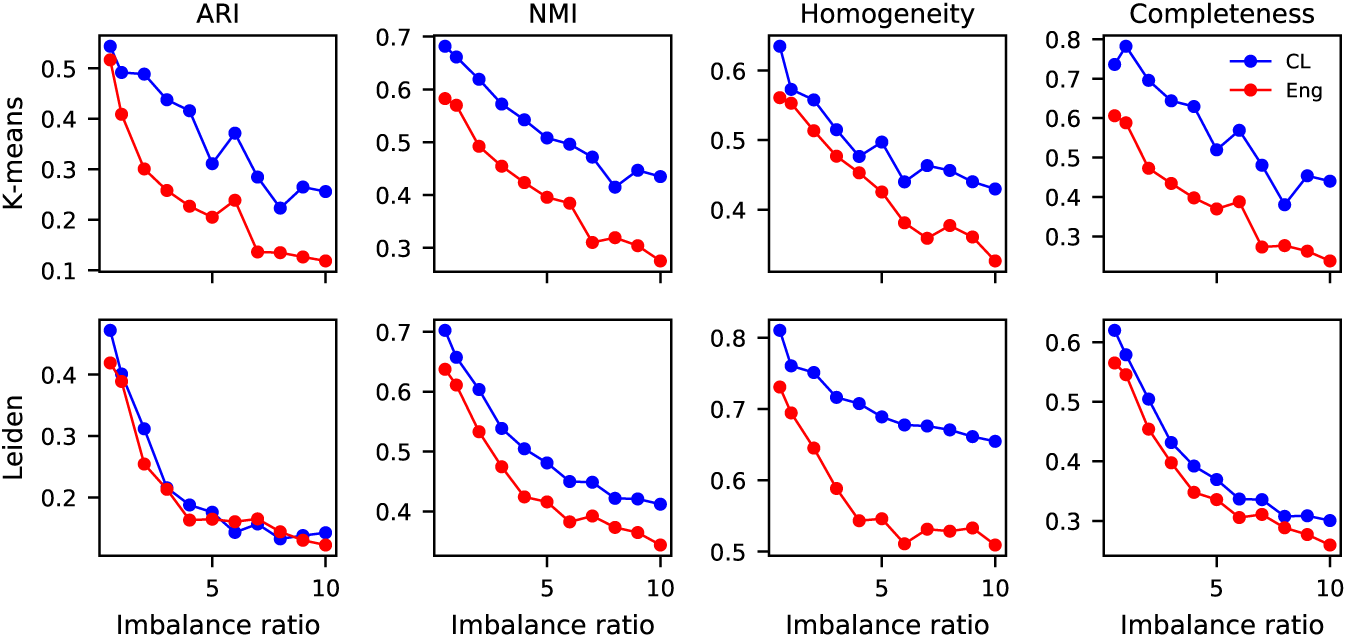
Learned features improve clustering performance on imbalanced cell phenotype data. Clustering performance of learned (CL) and engineered (Eng) feature representations was evaluated using two unsupervised algorithms—K-means and Leiden community detection—on a ground truth dataset of cell phenotypes. Experiments were conducted across a range of data imbalance ratios, defined as the ratio of immune cells to rare cells, from 0.5 to 10. Clustering quality was assessed using adjusted Rand index (ARI), normalized mutual information (NMI), homogeneity, and completeness scores.

Across all imbalance ratios, learned features demonstrated improved clustering performance over engineered features. For example, with K-means clustering at the lowest imbalance ratio, learned features (CL) achieved Completeness value of 0.74 compared to 0.61 for engineered features. Similar trends were observed for NMI (0.68 versus 0.58), Homogeneity (0.63 versus 0.56), and ARI (0.54 versus 0.52). As the imbalance ratio increased, performance declined across all methods; however, learned features maintained a performance margin of 5%–25% over engineered features, particularly in Homogeneity and NMI metrics. Aside from similar performance regarding ARI score, this advantage persisted with Leiden clustering, where learned features yielded a 8%-14% improvement in all other metrics at lowest ratios and retained higher performance across increasing imbalance ratio.

### 2.6 Learned features enable accurate enumeration of rare cell phenotypes in WSI

Having established the utility of learned features compared to traditional features, we next sought to evaluate their performance in a common downstream application. Accurate enumeration of rare tumor-associated cell phenotypes is a critical application of cell-based LBx, particularly for clinical decision-making and biomarker assessment. However, reliable identification remains challenging due to the extreme rarity of these cells and potential presence of technical artifacts and non-tumor rare events in WSI data. We evaluated the efficiency of our learned feature representations in detecting distinct rare cell phenotypes in two different experiments: (1) spike-in experiments with contrived samples of known cell lines at defined rarity, and (2) patient samples containing two different CTC phenotypes, canonical epithelial phenotype (CTCs) as well as immune-imCTCs.

In the first setting, two contrived WSIs containing either SK-BR-3 or HPAEC cells were used to train a multilayer perceptron classifier on learned features to classify each single cell as SK-BR-3, HPAEC, or other. We evaluated this model on 7 additional WSIs composed of various combinations of these cell lines and controls, totaling over 15 million single-cell events, including 1,902 SK-BR-3 cells and 1,286 HPAEC cells. The classifier achieved high overall precision (0.886 for SK-BR-3, 0.971 for HPAEC) and recall (0.982 for SK-BR-3, 0.903 for HPAEC), with an area under the precision-recall curve (AUPRC) of 0.957 and 0.962 for SK-BR-3 and HPAEC, respectively (Figure 5a). Further, classification performance remained consistently high across different slides, with an average F1-score of greater than 0.93 on both cell lines (Figure 5b).

**Fig. 5.**
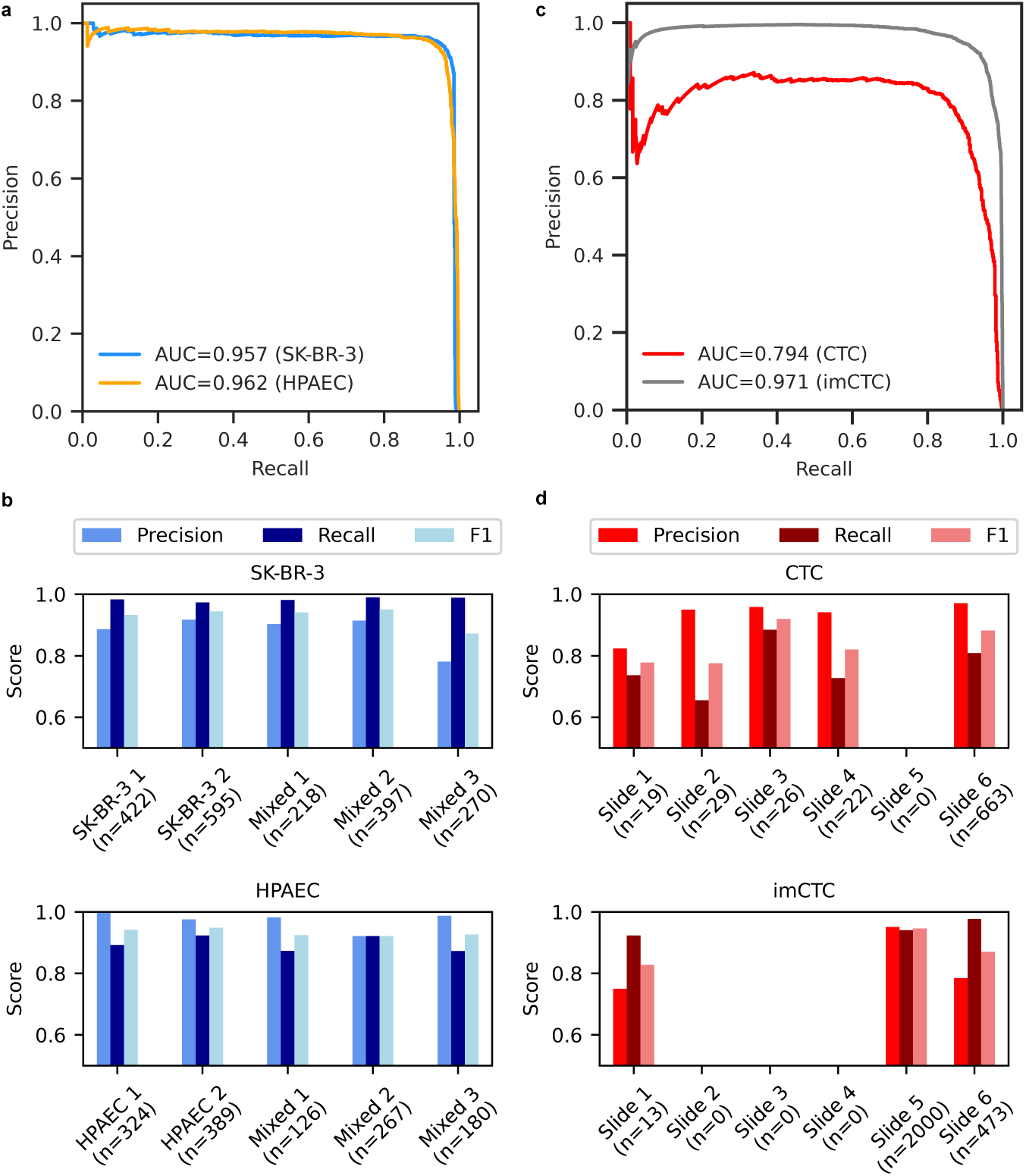
Learned features support accurate enumeration of rare cell phenotypes in both contrived and patient-derived WSI data. (a) Precision-recall curves for classification of SK-BR-3 and HPAEC cell lines in spike-in WSI data. (b) Per-slide performance metrics (precision, recall, F1-score) for SK-BR-3 and HPAEC classification across test WSIs containing different spike-in configurations. (c) Precision-recall curves for detection of CTC and immune-like CTC (imCTC) phenotypes in breast cancer patient WSIs. (d) Slide-wise performance metrics for CTC and imCTC classification across six held-out patient WSIs.

To assess its clinical utility, we applied the framework to a cohort of 8 samples from late-stage breast cancer patients with manually annotated CTCs and imCTCs. A classifier was trained on learned features extracted from two samples, and tested on six held-out samples comprising over 13 million cells. Despite significantly variable CTC frequencies across samples, from 19 to 663 CTCs and from 13 to 2000 imCTCs (representing 0.006% and 0.018% of total events), the model achieved overall precision values of 0.965 for CTCs and 0.912 for imCTCs, with recall values of 0.801 and 0.947, respectively. The AUPRCs were 0.794 for CTCs and 0.971 for imCTCs (Figure 5c). Slide-level analysis showed consistent performance, with average F1-score of 0.835 for CTCs and 0.881 for imCTCs (Figure 5d).

## 3 Discussion

Cell-based liquid biopsies hold a great promise for longitudinal monitoring and minimal residual disease identification in cancers. A variety of enrichment-based and enrichment-free approaches have revealed substantial heterogeneity and plasticity among rare, tumor-associated cells in circulation [13–15, 17–19]. Deconvoluting the heterogeneity of these circulating tumor-associated cells is feasible in enrichment-free platforms and can yield critical patient-specific insights and drive the discovery of novel biomarkers, ultimately improving patient outcomes [15]. Currently, analysis of distinct cell phenotypes is conducted through classical image processing methods using engineered cell image features, which are sensitive to technical variations, and often necessitate additional human annotation, which is prone to significant human bias.

Here, we developed a deep learning framework that leverages contrastive learning to develop features of single-cell LBx images to discriminate cellular populations in a robust and precise manner from unenriched LBx samples. Our results show that learned features are capable of accurately separating diverse tumor-associated phenotypes as well as immune cell subclasses. While previous work demonstrated comparable performance in supervised classification of 6 different cell types in EpCAM-enriched samples, their enrichment-based approach limited phenotypic diversity of single cells. By training and applying our feature encoding model on enrichment-free WSI data, we overcome this limitation and demonstrate strong performance in classification of 10 distinct classes, including 3 different CTC phenotypes. This enables the systematic study of multiple distinct tumor-associated phenotypes in parallel, along with distinct immune cell subpopulations that may be tumor-reactive, immunosuppressive, or otherwise indicative of tumor-associated signaling within the peripheral blood microenvironment. Such granularity is essential for capturing the full heterogeneity of circulating cells in liquid biopsy.

Our results confirmed that the learned features were more robust to regional blur, a common technical variation in WSI, than engineered features, suggesting that deep representations may help mitigate spurious rarity caused by imaging artifacts. This robustness is especially valuable in rare cell studies, where suboptimal image quality can significantly impact downstream analysis, including phenotyping classification, clustering, and outlier detection. Similar findings have been reported in other contexts where deep learning features demonstrated increased tolerance to image noise and blur compared to engineered features [45, 46].

Further, we demonstrate the capability of the learned features in outlier detection, a common task in the study of rare cells. The learned features yielded a higher concentration of rare cell lines among detected outliers in our contrived samples than engineered features. The consistently improved performance across multiple outlier detection methods suggests that the observed gains are attributable to the learned feature space rather than the specific detection algorithm. Moreover, the comparable detection performance of SK-BR-3 and HPAEC cells in the learned feature space contrasts with the biased detection favoring HPAECs when using engineered features. This indicates that the learned features promote more phenotype-agnostic performance, enabling balanced detection of rare cells across diverse phenotypes. Engineered feature space can, in principle, be selected for improved detection of each rare cell phenotype as well. However, this process requires prior knowledge of the target phenotypes, hindering the study of previously unknown rare cell phenotypes.

Moreover, learned features show significant improvements in the capacity to cluster cell phenotypes over engineered features in the presence of data imbalance, a natural property of cell phenotype populations in WSI data. Given the sensitivity of clustering algorithms to class imbalance and the ongoing development of specialized methods to address this limitation, our framework offers a robust feature space, as an orthogonal improvement, that enhances the ability to identify and discover novel cell phenotypes using current algorithms.

After establishing the utility of learned features in comparison to engineered features, our results revealed that the learned features enable precise identification and enumeration of two different target cell lines, mimicking CTCs (SK-BR-3) and CECs (HPAEC). Our work on enrichment-free samples resulted in F1-scores of 0.934 and 0.937 for SK-BR-3 and HPAEC, respectively, achieving equivalent performance to size-based enrichment studies that leverage supervised classification directly on cell images in the enriched cell population [47]. These results demonstrate that size-based microfiltration might be excessive for robust identification of circulating rare cell phenotypes. Moreover, relatively consistent performance across independent samples in our results demonstrate that the features we provide enable reproducible classification and enumeration of target cell phenotypes, offering reliable application of features in enumeration-based studies of cell biomarkers in patient cohorts.

Finally, we extended our classification analysis to patient samples by demonstrating that the learned feature representations enabled accurate identification and enumeration of two phenotypically distinct circulating tumor cell CTC populations. Although the model was trained on data from only two patients, it achieved average F1-scores of 0.835 and 0.881 for CTCs and imCTCs, respectively, when evaluated on independent patient samples. This strong performance, despite limited training data, highlights the capacity of feature space to generalize to unforeseen patient samples. A recently published work proposes a supervised model which has been trained on EpCAM-enriched patient samples, achieving an F1-score of 0.886 [48] with a limited detection capacity to only canonical epithelial CTCs. While having comparable performance scores, our framework analyzes enrichment-free WSI data and allows for the identification of a broader spectrum of multiple tumor-associated cell phenotypes. Together with its high performance, these results highlight the potential of our features for a broad clinical applicability. To the best of our knowledge, this is the first application of deep learning–derived features for simultaneous enumeration of multiple CTC phenotypes from patient data. While further validation across larger and more diverse cohorts is warranted, these findings suggest a promising potential for scaling this framework in support of rare cell phenotype enumeration and classification in translational studies.

One of the limitations of this work is that the feature encoding model was trained on a small number of patient samples, limiting the generalizability of the existing feature space. Inclusion of a larger, more diverse patient population across multiple cancer types and stages in training, will improve the performance of our general-purpose LBx feature extraction. Further, the limited training data of only two patient samples for CTC enumeration hinders the performance of classification models across novel patients samples. Given the inter- and intra-patients variability of CTCs and heterogeneity of cells other than CTCs [49], including more samples in the training data can significantly improve the accuracy of classification models in identifying the target cells.

Taken together, our findings demonstrate that contrastive learning yields single-cell IF image representations capable of facilitating robust phenotype classification, outlier detection, cell clustering, and supervised enumeration of circulating tumor-associated cells, even under realistic imaging artifacts and severe class imbalance. By removing the dependency on experimental enrichment and manual curation, this framework enables high-throughput pipelines for circulating rare-cell analysis that are more scalable, reproducible, and efficient than current practice. The same feature space can also be aggregated in the future with methodologies such as multiple-instance learning to learn patient-level biomarkers, paralleling recent advances in histopathology. Overall, these results position deep phenotyping of single-cell, enrichment-free, liquid-biopsy images as a powerful, generalizable approach for cancer monitoring and biomarker discovery.

## 4 Methods

### 4.1 Patient cohorts

Patient samples included in this study were assembled from multiple, previously published independent cohorts spanning several IRB-approved clinical studies. A collection of WSI data were selected from deidentified patient samples to provide a diverse population of rare events for training the segmentation and feature extraction modules as well as to cover the heterogeneity in immune populations for training the immune cell classifier used for depleting leukocytes to curate the training data for the feature extraction module. All samples were collected under protocols approved by the respective institutional review boards (IRBs), and written informed consent was obtained from all participants in accordance with the Declaration of Helsinki. Prior to analysis, all samples and associated metadata were de-identified to ensure patient privacy. Cohort-specific details, including cancer type/condition, study reference, number of samples used in this study, and IRB protocol number are summarized in Table 3.

### 4.2 Sample collection and preparation

LBx samples, predominantly peripheral blood, ( 8 mL) were collected in 10 mL tubes (Streck, Cell-free DNA BCT) at the clinical site and processed within 48 hours of collection [52]. Upon receiving the samples, cell counts were measured using Hematology Analyzer to determine the volume of blood needed for each slide to plate about 3 million cells per slide. In processing, isotonic ammonium chloride (A649-3, Fisher Scientific) erythrocyte lysis was conducted on each sample and the remaining cellular fraction is was plated on custom-made cell adhesive glass slides (Marienfeld, Lauda, Germany), generating between 8 and 16 slides per blood specimen, depending on the cellular density of the sample. Cells were plated on each slide, incubated with 7% BSA, dried, and stored at -80° C for future studies [50–52].

### 4.3 Immunofluorescence staining protocol

Slides were thawed for one hour and stained using the IntelliPATH FLX autostainer (Biocare Medical LLC) using a 4-color staining protocol. Each slide was first fixed with 2% PFA (stock 16%, 15710, Electron Microscopy Sciences) in 1x TBS (TWB945, Biocare Medical LLC) for 20 minutes. Then, the slides were incubated for 4 hours with 2.5 *µ*g/mL of a mouse IgG1 anti-human CD31 monoclonal antibody conjugated to AlexaFluor® 647 (clone: WM59, MCA1738A647, BioRad) and 100 *µ*g/mL of goat anti-mouse IgG monoclonal Fab fragments (115–007–003, Jackson ImmunoResearch). Cells were then permeabilized for 5 minutes using 100% methanol (A412-1, Fisher Scientific) and blocked for 20 minutes with 10% filtered goat serum (S26-LITER, Emd Millipore) in TBS to prevent non-specific interaction of primary and secondary antibodies. Goat serum was used as an antibody dilution buffer to the following steps of the staining protocol. Slides were incubated for 2 hours at room temperature with a primary antibody cocktail including mouse IgG1/IgG2a anti-human pan-cytokeratin monoclonal antibodies (CK1, CK4, CK5, CK6, CK8, CK10, CK13, CK18) (C2562-.5ML, Sigma-Aldrich), and CK19 antibody (M088801-2, Dako North American), rabbit IgG anti-human vimentin monoclonal antibody conjugated to AlexaFluor® 488 (VIM, 9854BC, Cell Signaling Technology), and mouse IgG2a anti-human CD45 monoclonal antibody conjugated to AlexaFluor® 647 (CD45, MCA87A647, AbD Serotec). Finally, slides were incubated with goat anti-mouse IgG1 AlexaFluor® 555 (A21127, Thermo Fisher) secondary antibody and 4 ’6 diamidino-2-phenylindole (DAPI, D1306, Thermo Fisher), mounted in a glycerol-based mounting media (BP229-1, H5123, ICN10274750, Fischer Scientific), and cover-slipped (12545J, Fisher Scientific) [15].

### 4.4 Image acquisition

Stained slides were then imaged using a in-house automated fluorescence scanning microscope with 10x objective and 100x overall magnification, yielding 2304 frames per slide for each of the four immunofluorescence channels [50]. Collected images were stored as 16-bit TIFF format (13621004). The total image size for one slide is approximately 16 GBs.

### 4.5 Engineered feature extraction

The engineered features for all analyses were extracted using the EBImage package (version 4.48) in R (version 4.4.2) provided the image and the cell mask. The 368 features included cell size, eccentricity, haralick texture features and intensity statistics including mean, median, standard deviation and different quantiles from each individual channel image as well as combination of channel pairs.

### 4.6 Segmentation model

#### 4.6.1 Dataset

As a ground truth dataset for segmentation, a set of 634 (256 x 256 pixels) images were collected as crops from frame images of a subset of previously-published rare cell phenotype dataset [52, 53]. These images were selected such that they include a diverse set of cell phenotypes to ensure they represent the heterogeneity across cellular morphologies for improved masking. After running cyto3, the pre-trained Cellpose model, on these images, the generated masks were manually corrected using the Cellpose graphical user interface (version 3.0.11).

#### 4.6.2 Architecture, training and evaluation

We proceeded to train a custom segmentation model using the Cellpose backbone architecture, with no pretrained weights [42]. To evaluate the performance of our custom-trained Cellpose model, we generated a dataset of 153 unseen 256 x 256 pixel test crops that were manually masked using the Cellpose Graphical Interface [42]. To evaluate instance segmentation performance, we computed the object-level F1-score at multiple intersection-over-union (IoU) thresholds [54]. This metric quantifies the accuracy of predicted object masks by comparing them to ground truth annotations based on their spatial overlap. The IoU is defined as the ratio between the area of intersection and the area of union of a predicted and a ground truth object mask. A predicted object is considered a true positive (TP) if its IoU with a ground truth object exceeds a specified threshold, and each ground truth object can be matched to at most one prediction. Unmatched predictions are counted as false positives (FP), and unmatched ground truth objects as false negatives (FN). For each IoU threshold, F1-score is calculated by Equation 1. This evaluation captures both the detection and localization accuracy of instance-level predictions, with increasing IoU thresholds enforcing stricter spatial agreement for a match.

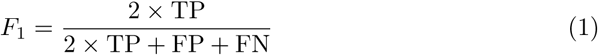

### 4.7 Leukocyte classifier architecture and training

#### 4.7.1 Dataset

We collected a dataset of 18,298 annotated cells from previously published datasets in breast cancer [52], prostate cancer [15], multiple myeloma [53], bladder cancer [**?**], colorectal cancer [55], non-small cell lung cancer [56], upper tract urothelial carcinoma [57] and non-cancerous post-acute covid syndrome [58]. The dataset was annotated in the following classes: leukocyte and non-leukocyte. The dataset was split 80% train, 20% validation. The balance of the training data was 7,319 leukocytes to 7,319 non-leukocytes. The validation dataset contained 3,660 total cells, with 1,830 of those annotated as leukocytes. Each cell image consisted of a centered, masked 75 x 75 image, where the mask was multiplied across the four immunofluorescent channels and concatenated as a fifth channel.

#### 4.7.2 Architecture, training and evaluation

The Leukocyte Classifier was trained using a PyTorch implementation of a CNN. The model consisted of four batch-normalized convolutional layers, activated by ReLU, each followed by a Max-Pooling layer of size (2,2), and a 2-layer dense network to reduce the features to class logits. The cross-entropy loss function was used to train the model. A learning rate of 0.0001 was used with the Adam optimizer to optimize the training. The model was trained for 25 epochs on one RTX4090 GPU, monitoring train and validation loss curves. AUROC and PR-ROC curves were generated on the validation data.

### 4.8 Feature encoding model

#### 4.8.1 Dataset

To train the representation learning model, we first negatively enriched the leukocyte population on 25 whole-slide images using the above Leukocyte Classifier. To ensure the model learned the rare-event space and the leukocyte space, we sub-sampled 450 leukocytes from each whole slide at random and added them to the collected non-leukocytes. The 25 instances of WSI data included breast, prostate, bladder, multiple myeloma, and non-cancerous post-acute covid syndrome samples. Of the 25 samples, 2 samples were taken from bone-marrow aspirates (one prostate cancer and one multiple myeloma), and 23 samples were taken from peripheral blood. After applying the Leukocyte Classifier, we were left with a total dataset of 117,821 non-leukocytes. Additionally, we randomly subsampled 450 predicted leukocytes from each slide, yielding a total dataset of 129,071 cells. The cells were split 80% training / 20% validation to yield 103,245 training events and 25,826 validation events. Each event consisted of a 75 x 75 centered, masked cell with the mask multiplied across the 4 immunofluorescent channels, as well as appended as a fifth channel.

#### 4.8.2 Architecture, training and evaluation

The architecture of the representation learning model comprises a CNN encoder architecture and a two-layer projection network. The CNN contains four convolutional layers, each followed by batch normalization. After the last convolutional layer a MaxPooling Layer with a 2x2 kernel, with stride of 2 is applied, followed by an Adaptive Pooling Layer with a 1x1 kernel. ReLU is used as the activation function following all convolutions. The features are then flattened before being passed to the two-layer projection network, where the loss is calculated and back-propagated.

For every minibatch, we use the normalized temperature-scaled cross entropy loss function [59]. In a given batch with *n* data points, each data point *x* is augmented twice to create *x_i_* and *x_j_*. The positive pair is forward propagated through the CNN encoder to extract representations *h_i_* and *h_j_*. These representation vectors are passed through the projection head to yield *z_i_*and *z_j_*. This loss function is represented in Equation 2.

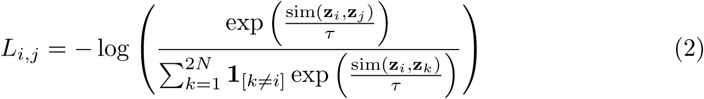

Although there is no negative sample mining, every sample in the training batch, aside from the positive pair, is considered a negative sample. The dot product between two data points gives the cosine similarity, sim(*z_i_*, *z_j_*). is the temperature parameter. The loss function aims to update the model weights such that the model maximizes the cosine similarity between the representations of the positive pair while minimizing the similarity between the positive and negative samples.

The training procedure is as recommended by SimCLR [38]. We selected hyperparameters of learning rate, temperature, learning-rate scheduler, latent-dimension size, projection-dimension size, batch size, weight decay, and training epochs using random search. The final model was trained using the LambdaLR learning rate scheduler, with a starting learning rate of 0.001 and a linear warm up period of 10 epochs to reach a maximum learning rate of 0.01. Temperature was set to 0.13. The representation dimension, h, was a 128-dimensional vector while the projection dimension was 64. Batch size was set to 1024. Weight decay was set to 0.0001. The model was trained for 50 epochs on one RTX4090 GPU.

The choice and configuration of image augmentations play a critical role in contrastive learning. In our approach, we apply a series of random transformations to each image in a pair, independently and in a fixed order, with each augmentation having its own probability p of being applied. First, we apply a color jitter on all channels (brightness=0.4, contrast=0.4, saturation=0.4, hue=0.2, p=0.5). We then apply a random rotation between -180° and 180° (p=0.5), followed by random horizontal and vertical flips (both p=0.5). Next, an affine translation of up to 15 pixels off-center in both the x and y directions is added (p=0.5), after which we apply a Gaussian blur (p=0.5) using a 3-pixel kernel and a sigma chosen randomly between 0.5 and 3.0. Finally, we randomly crop the image to between 50% and 100% of its original size, then resize it back to 75 × 75 (p=0.5).

These parameters were selected through careful visual evaluation of their effects on immunofluorescent images, ensuring that we preserved crucial fluorescent signals while encouraging the model to generalize across a wide range of cellular phenotypes.

### 4.9 Cell phenotype classification model

#### 4.9.1 Dataset

Cells previously characterized through IMC for highly multiplexed proteomics were collected for evaluating the feature encoding model [13, 16, 43]. This dataset was further expanded by adding phenotypically matched cells collected by trained analysts. The cells evaluated here were independent of those present in the representation learning training. The IMC analysis was conducted on a CyTOF Helios imaging mass cytometer (Fluidigm). A table of antibody clones used in these studies and their associated cell type can be found in Supplementary Table 3. Antibodies not directly conjugated to metals were conjugated in the lab with a Maxpar antibody labeling kit. Single cells were segmented through a trained ilastik (version 1.3.3) model and ion count per cell was extracted using CellProfiler (version 3.15) in a pipeline developed by the Bodenmiller lab [60]. The total number of cells previously annotated was 2,166 and the cells spanned seven classes: immune-like CTCs (imCTC), canonical epithelial CTCs (CTC), platelet-coated CTCs (pcCTC), endothelial cells (CEC), megakaryocytes, fibroblasts, and cells with large nuclei that expressed no other markers (Large Nuclei). Thresholds for each marker were defined by observing the ion count histograms. The 2,166 rare cells were split 80% train, 20% validation.

#### 4.9.2 Architecture, training and evaluation

A logistic regression model was trained on the 80% split training data from scikit-learn package (version 1.5.2). The model was trained for a maximum of 10,000 iterations until convergence. Accuracy, precision, recall, and f1-score were calculated for each class on the 20% held out validation dataset.

### 4.10 Sensitivity to blur analysis

#### 4.10.1 Dataset

The dataset utilized in this experiment is the same as described in 4.9.

#### 4.10.2 Analysis

For each cell present in our dataset, we generated features using our contrastive learning feature set and a set of engineered cell features on sharp images and those with synthetically generated blurring using gaussian blur with fixed, increasing values of sigma (0.1, 0.2, 0.3, 0.5, 0.75, 1, 1.25, 1.5). We evaluate the pairwise cosine distance between the features for the sharp cell image and its associated blurred image to evaluate the change in the distance between the two vectors.

### 4.11 Outlier detection analysis

#### 4.11.1 Dataset

We generated control WSI data by adding, at a known concentration of 1 cell-line cell to 10,000 leukocytes, SK-BR-3, HPAEC, or both cell lines to tubes of blood collected from healthy donors. The slides were processed in the same manner as is described above (Sample Collection and Preparation, Immunofluorescence staining protocol). Four slides were generated from each tube of blood collected from a healthy donor: one containing no cell lines, one containing SK-BR-3 cells only, one containing HPAEC cells only, and one containing both SK-BR-3 and HPAEC cells. We replicated this process for three healthy donors, yielding a total of 9 control slides. Each instance was processed through the Cellpose segmentation and the representation learning model to extract 128-dimensional feature vectors for each event.

SK-BR-3 cells and HPAEC cells were manually annotated from each WSI data using annotateEZ, an annotation tool developed in-house (see code availability section). SK-BR-3 cells were selected for their large size, expression of cytokeratin, and negativity for vimentin and CD45/CD31. HPAEC cells were selected for their expression of CD45/CD31 (multiplexed in one channel), vimentin, and large size. We have previously shown that HPAEC cells have variable expression levels of cytokeratin [43]. Across all slides, 2,641 cells were classified as SK-BR-3, and 1,691 cells were classified as HPAEC.

#### 4.11.2 Analysis

We evaluated the performance of features extracted through contrastive learning in detecting true outliers and compared the performance to engineered features. The contrastively learned features have 128 dimensions, whereas the engineered features have 368 dimensions (see engineered feature extraction). For three outlier detection algorithms (COPOD, ECOD, iForest), we evaluated the area under the ROC curve for the first 2,500 events (representing approximately 0.1% of the data in a WSI). All outlier detection algorithms were implemented using pyOD cite pyOD. For all outlier detection algorithms, the level of contamination was set to 0.001. The number of estimators used in iForest was 100.

### 4.12 Clustering analysis

#### 4.12.1 Dataset

This experiment utilized the same dataset as described in 4.8. We further sampled leukocytes (labeled as either monocytes, granulocytes, or lymphocytes) to change the imbalance ratio between leukocytes and rare cell classes.

#### 4.12.2 Analysis

We evaluated clustering performance using two unsupervised algorithms: K-means and Leiden community detection. For K-means, clustering was performed using the KMeans implementation from scikit-learn with the number of clusters set to 10 and a fixed random seed to ensure reproducibility. For Leiden clustering, a k-nearest neighbors (k-NN) graph was constructed using cosine similarity with neighborhood size set to 15. The resulting graph was symmetrized and passed to the Leiden algorithm (via the leidenalg package), using the modularity vertex partition scheme. Clustering results from both methods were evaluated against ground truth labels using standard metrics (Adjusted Rand Index, Normalized Mutual Information, Homogeneity, Completeness), as described in the clustering evaluation section.

### 4.13 Contrived cell phenotype identification and classification model

#### 4.13.1 Dataset

This experiment utilized the same dataset as described in 4.11.

#### 4.13.2 Architecture, training and evaluation

2 of 9 slides, one containing SK-BR-3 cells only and another containing HPAEC cells only, were used to train a multiclass classifier to evaluate the ability of the embeddings to learn cell-type on a limited set of data. The multiclass classifier consisted of a two-layer fully connected multilayer perceptron model. The model was trained for 100 epochs with a learning rate of 0.01 and optimized with the Adam optimizer. The training data consisted of 6,636 events classified as others, 683 events classified as SK-BR-3, and 408 events classified as HPAEC, which was split 80% training and 20% validation. Loss was monitored for both the training and validation sets during training. We evaluated the performance of the representation-learning model by applying the classifier across the 7 remaining independent slides. For each slide, we calculated the precision, recall, and F1-score. We additionally aggregated all of the slides and calculated overall performance metrics.

### 4.14 Patient cell phenotype identification and classification model

#### 4.14.1 Dataset

We collected 8 previously characterized WSIs from patients with metastatic breast cancer. These patients have previously been characterized as having CTCs, imCTCs, or both cell phenotypes in their samples. The slides were processed in the same manner as is described above (Sample Collection and Preparation, Immunofluorescence staining protocol). Each instance was processed through the Cellpose segmentation and the representation learning model to extract 128-dimensional feature vectors for each event. CTCs and imCTCs were manually annotated from each WSI data using annotateEZ. CTCs were selected for their large size, expression of cytokeratin, and negativity for vimentin and CD45/CD31. imCTCs were selected for their expression of cytokeratin, CD45/CD31 (multiplexed in one channel), and large size. Across all eight slides, 936 cells were classified as CTCs, and 5,220 cells were classified as imCTC.

#### 4.14.2 Architecture, training and evaluation

2 of 8 slides were used to train a multiclass classifier to evaluate the ability of the embeddings to learn cell-type on a limited set of data. The multiclass classifier consisted of a three-layer fully connected multilayer perceptron model. The model was trained for 50 epochs with a learning rate of 0.001, a weight decay of 0.0001 and optimized with the Adam optimizer. The training data consisted of 5,000 events classified as others, 177 events classified as CTCs, and 2734 events classified as imCTCs. We evaluated the performance of the representation-learning model by applying the classifier across the 6 remaining test slides. For each slide, we calculated the precision, recall, and F1-score, in addition to overall performance metrics.

## 5 Data Availability

The datasets generated and used for training and fine-tuning the models have been deposited in the BloodPAC Data Commons and are available under accession ID BPDC000147.

## 6 Code Availability

All the codes developed for the model training and analysis are available at https://github.com/CSI-Cancer/deep_phenotyping. The image annotation tool, AnnotateEZ, is available to the public at https://github.com/aminnaghdloo/annotateEZ.

## Acknowledgements

The work was funded in whole or in part by NIH NCI U01CA258013 (P.K., J.H.), Department of Defense CA220785 (P.K., J.H.), Breast Cancer Research Foundation BCRF-24-089 (P.K., J.H.), Dr. Miriam and Sheldon G. Adelson Medical Research Foundation (P.K., J.H.), Ming Hsieh Institute for Research on Engineering-Medicine for Cancer (P.K., A.O.), and NCI’s USC Norris Comprehensive Cancer Center (CORE) Support P30CA014089 (P.K., J.H.). This work also received institutional support from the USC Michelson Center Convergent Science Institute in Cancer, Winnie and James Hart Endowed Fellowship (A.N.), Schlegel Family Endowed Fellowship (D.T.), and Vassiliadis Research Fund. The content is solely the responsibility of the authors and does not necessarily represent the official views of the National Institutes of Health. Further, we thank the patients and their caregivers who consented to this study. We also thank the clinical research staff who contributed to the study. We are grateful to past and current technical staff at CSI-Cancer for processing the samples.

## Supplementary information

The supplementary figures and data are included in the accompanied supplementary file.

## Author contributions

These authors contributed equally: Amin Naghdloo, Dean Tessone, and Rajiv Mandya Nagaraju. Conceptualization, A.N., D.T., R.M.N., and P.K.; methodology, A.N., D.T., R.M.N, and B.Z.; validation, A.N. and D.T.; formal analysis, A.N. and D.T.; resources, J.H and P.K.; data curation A.N., D.T., and S.L., writing, A.N., D.T., B.Z., writing review, all authors; visualizations, A.N. and D.T.; code authorship, A.N., D.T., R.M.N, B.Z., J.K., and S.L.; code orchestration, D.T., R.M.N, and B.Z., supervision, A.O., J.H., and P.K., project administration, P.K.; funding acquisition, J.H., and P.K.;

