## Supplementary Figures and Tables for "Contrastive Representation Learning for Single Cell Phenotyping in Whole Slide Imaging of Enrichment-free Liquid Biopsy"

### Supplementary Material

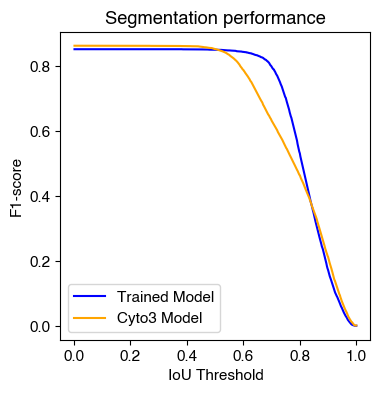

**Supplementary Figure 1. Performance curve of retrained segmentation model compared to general-purpose Cyto3 model provided by Cellpose.** Object-level F1-score is plotted as a function of IoU threshold for both the retrained model (blue) and the Cyto3 model, which is provided as the pretrained all-purpose model by Cellpose (orange). The retrained model consistently outperforms the baseline across a broad range of thresholds, indicating successful adaptation of the segmentation model to LBx WSI data.

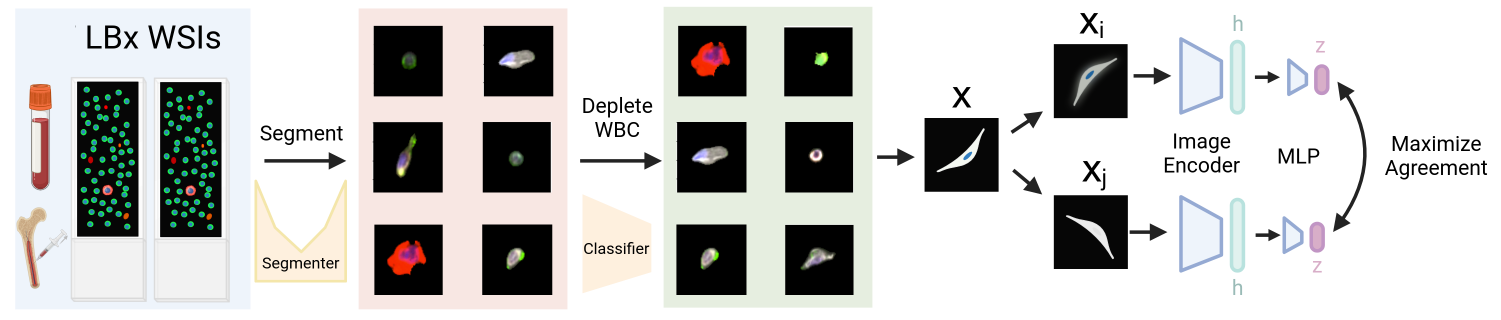

**Supplementary Figure 2. Overview of training feature encoding module.** The feature encoding module was trained on cells sub-sampled from 25 WSIs after segmentation. The cells are passed through a contrastively-trained feature encoder. The features are then evaluated in three downstream tasks. 1) Assessing the ability of the extracted features to describe tumor-associated cellular populations. 2) Assessing the ability of the extracted features to describe the immune population. 3) Assessing the ability of the extracted features to automatically analyze WSIs for tumor-derived analytes.

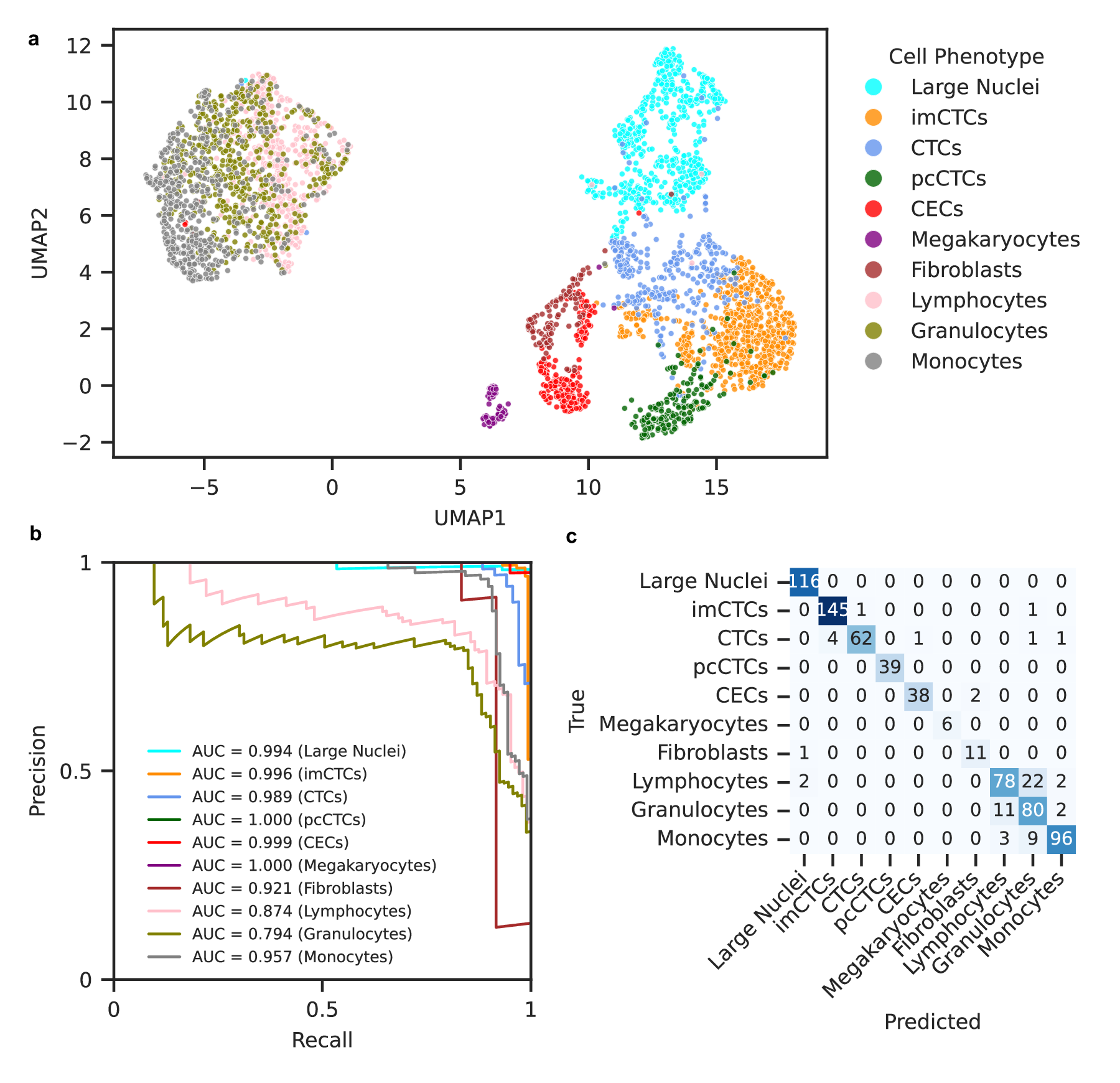

**Supplementary Figure 3. Engineered feature representations stratify diverse single-cell phenotypes.** Engineered features were evaluated on the same ground truth of cell phenotypes. **a.** UMAP projection of single cells based on engineered features, colored by ground truth phenotype labels. **b.** Precision-recall curves for each phenotype on the held-out test set, with areas under the curve (AUCs) shown in the legend. **c.** Confusion matrix of the logistic regression multi-class classifier evaluated on the test data.

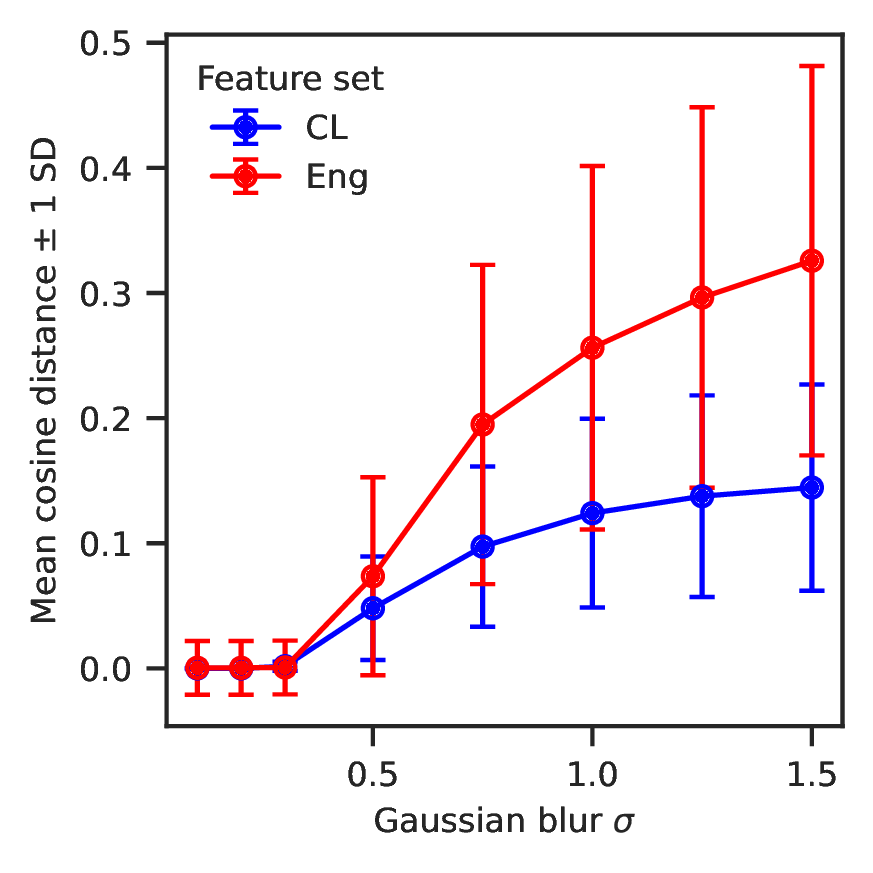

**Supplementary Figure 4. Learned features by our framework are more robust to blur.** Sensitivity to blur as a common technical artifact in whole slide imaging is evaluated for learned features via contrastive learning (CL), engineered features (Eng). Average cosine distance between original and blurred image features is used as a metric to quantify the effect of blur on feature space. Engineering features are more impacted by blur than learned features.

**Supplementary Table 1. Annotated rare cells and leukocytes used to train the leukocyte classifier and segmentation model.** Previously published annotated datasets were used to train the leukocyte classifier and segmentation model. Total number of patients in each cancer type from which that data was derived is presented here.

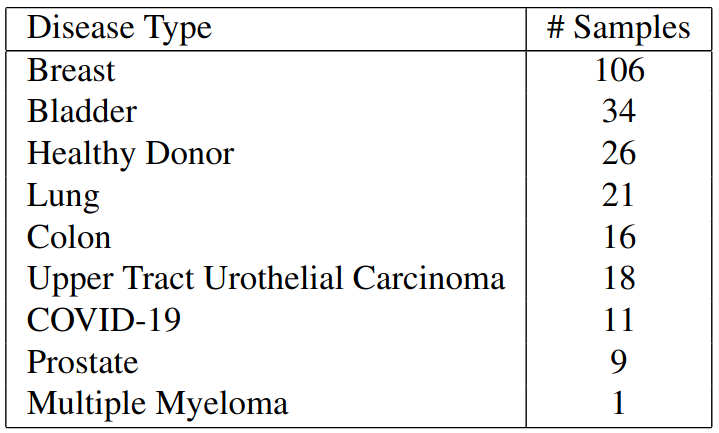

**Supplementary Table 2. WSI sample distribution by disease type and Sample type used to train feature encoding Model.** Whole slide image data were collected from 25 samples from a variety of disease types and were used to train the feature encoding model. The number of WSIs from each disease type and whether they were peripheral blood or bone marrow is presented here.

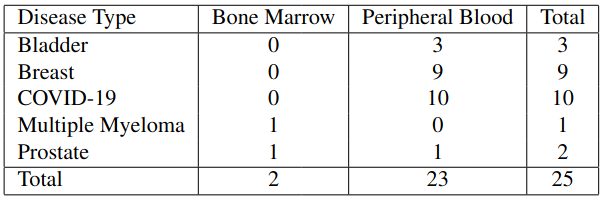

**Supplementary Table 3. Antibodies Clones used in IMC experiments for Cell Phenotyping.** Antibodies are labeled by their clone and catalogue number. The cell type associated with each antibody level positivity is labeled as well.

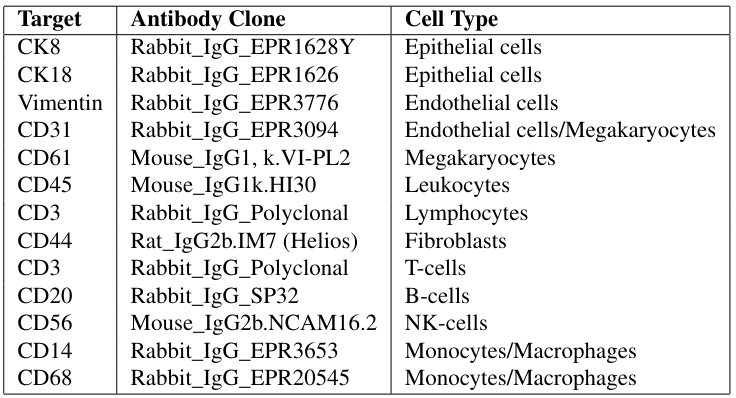
